# Discovery of new and variant *Potyviruses* in *Asparagales* plants from a Dutch urban botanic garden

**DOI:** 10.1101/2024.10.09.617369

**Authors:** Rob J. Dekker, Han Rauwerda, Wim C. de Leeuw, Marina van Olst, Wim A. Ensink, Selina van Leeuwen, Claartje Meijs, Moezammin M.M. Baksi, Timo M. Breit, Martijs J. Jonker

## Abstract

Although viruses play an important role in human health and plant health, most viruses are still undetected. One of the reasons is that they may evolve in distinct natural habitats that are not yet extensively sampled for virus analysis. An example of such a habitat is urban botanic gardens. We analyzed 25 *Asparagales* plants with a mild to severe disease phenotype from a Dutch urban botanic garden for the presence of known an unknown virus types and virus variants by smallRNA-seq and RNA-seq. We found in all samples evidence for (past) *Potyviridae* presence, mostly Ornithogalum virus (OV) and Ornithogalum mosaic virus (OrMV), as well as a new Iris mild mosaic virus variant (IMMV) and a yet unknown species of Potyvirus. Also, presence of a new Betaflexivirus, a new Polerovirus and a new Phenuivirus were detected. Most analyzed plants harbored multiple viruses, 18 out of 25 plants showed evidence for three to seven viruses and 12 out of 13 viruses were present in four to 11 samples. In this study, we describe the characteristics of a newly discovered Potyvirus and identify several variants of known potyviruses. We place these findings in the context of known viruses. However, we were unable to link these potyviruses to any specific disease phenotype.

## Introduction

With the introduction of affordable metagenomics approaches, it has become rapidly clear that only a miniscule proportion of the total virosphere is known (Zhang *et al*., 2018, Paez-Espino *et al*., 2016; Neri *et al*., 2022). Since many viruses are pathogenic for their host, viruses are an important factor in mammalian and plant health and thus human health and food security. Also, viruses often show us remarkable RNA characteristics that are very important in understanding RNA biology (Scholthof and Scholthof, 2023). Hence, it is crucial to expand our knowledge on viruses. Detection is challenging because many viruses cannot be easily identified using standard sequence homology-based methods, often referred to as viral RNA dark matter (Krishnamurthy and Wang, 2017). Additionally, numerous viruses reside in hosts that have not yet been screened, or they inhabit host plants of screened species that live in specific environments.

One of these scenarios may apply to plants in urban botanic gardens. Despite the potential for botanic gardens to serve as breeding ground for plant viruses, there is limited research available on virus presence in botanic gardens. We hypothesized that given the defined space of an urban botanic garden, the exchange policy of rare plant species between botanic gardens, plus the fact that most plants are kept for countless generations could result in an abundant presence of many specific viruses in urban botanic-garden plants. To investigate this, we screened plants from an urban botanic garden for known and unknown viruses by smallRNA-seq (sRNA-seq) and RNA-seq. RNA-seq would reveal the viruses currently present in a plant, while sRNA-seq, through virus-related siRNA, could indicate present, as well as past virus presence.

We sampled 25 *Asparagales* plants from a Dutch urban botanic garden and found one new *Potyvirus* and several *Potyvirus* variants (this report), a new *Polerovirus* (Dekker *et al*., 2024a), a new *Betaflexivirus* (Dekker *et al*., 2024b), and a new *Phenuivirus* (Dekker *et al*., 2024c).

## Material and methods

### Samples

Samples of leaves from 25 *Asparagales* plants were collected from Hortus Botanicus, a botanic garden in Amsterdam, the Netherlands, on February 14, 2019. Details about the plant genera can be found in Supplemental Table ST1.

### RNA isolation

S-RNA was isolated by grinding a flash-frozen ±1 cm^2^leaf fragment to fine powder using mortar and pestle, dissolving the powder in QIAzol Lysis Reagent (Qiagen) and purifying the RNA using the miRNeasy Mini Kit (Qiagen). Separation of the total RNA in a small (<200 nt) and large (>200 nt) fraction, including DNase treatment of the large RNA isolates, was performed as described in the manufacturer’s instructions. The concentration of the RNA was determined using a NanoDrop ND-2000 (Thermo Fisher Scientific) and RNA quality was assessed using the 2200 TapeStation System with Agilent RNA ScreenTapes (Agilent Technologies).

### RNA-sequencing

Barcoded sRNA-seq and RNA-seq libraries were generated using a Small RNA-seq Library Prep Kit (Lexogen) and a TruSeq Stranded Total RNA with Ribo-Zero Plant kit (Illumina), respectively. The size distribution of the libraries with indexed adapters was assessed using a 2200 TapeStation System with Agilent D1000 ScreenTapes (Agilent Technologies). The sRNA-seq libraries from samples S01 to S12 and from samples S14-S26 were clustered and sequenced at 2×75 bp and 1×75 bp, on a NextSeq 550 System using a NextSeq 500/550 Mid Output Kit v2.5 or a NextSeq 500/550 High Output Kit v2.5 (75 cycles or 150 cycles; Illumina), respectively. RNA-seq libraries were clustered and sequenced at 2×150 bp on a NovaSeq 6000 System using the NovaSeq 6000 S4 Reagent Kit v1.5 (300 cycles; Illumina).

### Bioinformatics analyses

Sequencing reads were trimmed using trimmomatic v0.39 (Bolger *et al*., 2014) [parameters: LEADING:3; TRAILING:3; SLIDINGWINDOW:4:15; MINLEN:19]. Mapping of the trimmed reads to the NCBI virus database was performed using Bowtie2 v2.4.1 (Langmead *et al*., 2012). Contigs were assembled from sRNA-seq data using all trimmed reads as input for SPAdes De Novo Assembler (Prjibelski *et al*., 2020) with parameter settings: only-assembler mode, coverage depth cutoff 10, and kmer length 17, 18 and 21. Assembly of contigs from RNA-seq data was performed with default settings. Assembly of contigs from RNA-seq data was performed with default settings. Scanning of contig sequences for potential RdRp-like proteins was performed using PalmScan (Babaian and Edgar, 2022) and LucaProt (Hou *et al*., 2023).

## Results and discussion

To get a better understanding of the virus presence in botanic garden plants, we performed an RNA-seq experiment, as well as a sRNA-seq experiment on 25 *Asparagales* plants from 14 different species with no, mild or severe symptoms of disease, obtained from the urban botanic garden in Amsterdam (Supplemental Table 1). The sequencing yielded sufficient short and long sequencing reads for each sample to allow subsequent metagenomics analysis (Supplemental Table 1).

### A new *Potyvirus* species in *Lachenalia aloides*

During the metagenomics sequence analyses of our (s)RNA-seq data sets, we came across a several new viruses. Previously, we reported on a new Betaflexivirus (Ferraria Betaflexivirus 1, FerBfV-1, Dekker *et al*., 2024b), a new Phenuivirus (Lachenalia Phenuivirus isolate 1, LacPhV-1, Dekker *et al*., 2024c), and a new Asparagaceae Polerovirus 1 (AspPolV-1, Dekker *et al*., 2024a) (Supplemental Table ST2). Here we report on the discovery of a new single-stranded positive strand *Potyvirus*. The genomic RNA sequence is 9.485 bp long, coding for a polyprotein of 3.065 aa (Figure 1).

**Figure 1.**
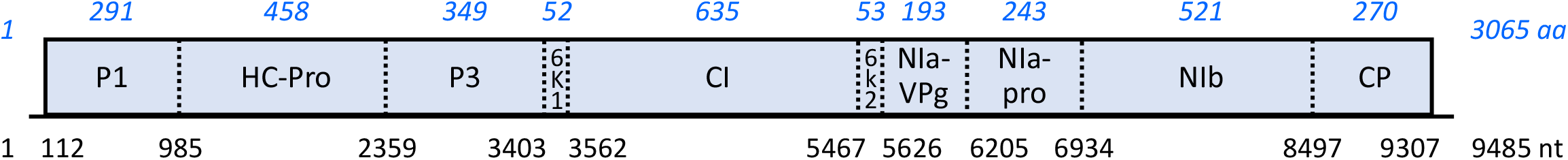
Schematic representation of the genome, the polyprotein, and the mature proteins of *Potyviridae*. Indicated are the positions of the proposed polyprotein cleavage sites (black) and mature protein sizes (blue) for Aloides Potyvirus-1 (AloPotV-1). The domains present in the mature proteins are; P - Protease; HC-Pro - Helper component protease; CI - Cylindrical inclusion; VPg - Viral protein genome-linked, NI - Nuclear inclusion, CP - Coat protein. NIb contains the RNA-Dependent-RNA-Polymerase (RdRp).

The virus with the closest similarity (RNA identity ∼60% and protein identity 53%) is Potyvirus Tuberose mild mosaic virus (GenBank: ON116187.1). These identity values are significantly lower than the Potyviridae virus species demarcation threshold (<76% identity in nucleotide sequence and <82% in in amnio acid sequence but exceeds the *Potyviridae* genus demarcation threshold of <46% identity in nucleotide sequence (Inoue-Nagata *et al*., 2022). Therefore, the new virus represents a new Potyvirus species. We were able to identify the signature *Potyvirus* polyprotein cleavage sites (Figure 1 and Table 2) and associated proteins, as well as a high-confidence RDRP palmID (score 65.1, Babaian and Edgar 2022, Supplemental Figure SF1). Upon mapping the (s)RNA-seq reads to the new Potyvirus sequence, we detected (s)RNA sequence evidence for this new Potyvirus in only one sample (S15, Supplemental Tables ST3 and ST4). As this sample originated from a *Lachenalia aloides* plant, we have named this new *Potyvirus* Aloides Potyvirus-1 (AloPotV-1, PQ213853).

### A known *Potyvirus* in *Albuca canadensis*

We observed also a known Potyvirus in four samples (S05, S10, S11, and S12, Supplemental Tables ST3 and ST4) as variants of the Iris mild mosaic virus (IMMV) (Table 1). The virus variant from sample (S11) with the most siRNA reads (IMMV-NL1-11, PQ156407, Supplemental Table 4) is 9.293 nucleotides long and contains an ORF encoding a 3.046 amino acid polyprotein. This variant shows 98.3% identity at the RNA level and 97.4% identity at the protein level to the closest matching Potyvirus sequence in GenBank (Iris mild mosaic virus isolate Ir, MN746770.1). Additionally, it exhibited virtually no differences at the polyprotein cleavage sites or RdRp motifs (Table 2, Supplemental Figure SF1).

### Reordering *Ornithogalum* viridae

To conclude, we identified several *Ornithogalum* virus variants (species *Potyvirus ornithogalitessellati*), all belonging to the family *Potyviridae* (Supplemental Tables ST3 and ST4). Trying to name these variant *Potyviruses*, we encountered issues with the existing nomenclature available for most, to this study relevant, *Potyviruses* in GenBank.

Applying reported species identity cut-off values (<76% identity in nucleotide sequence and <82% in amnio acid sequence for *Potyviridae* species demarcation (Inoue-Nagata *et al*., 2022), we identified several *Ornithogalum* virus variants and *Ornithogalum* mosaic virus variants that fall outside these demarcation values. However, similar virus genome sequences in GenBank were still classified as the same species, *Potyvirus ornithogalitessellati*. Therefore, the annotation of the Ornithogalum (mosaic) virus genomes in GenBank appears to not comply with the ICTV demarcation criteria. To address this issue, we compared all complete *Ornithogalum* virus sequences from GenBank, supplemented with the sequences from variants identified in this study (Supplemental Table ST5). This comparison revealed three distinct groups in an identity tree analysis, which we labelled *Ornithogalum virus 2* (OV-2), *Ornithogalum virus 3* (OV-3), *and Ornithogalum virus 6* (OV-6) (Figure 2A). Similarly, for *Ornithogalum mosaic virus* (Supplemental Table ST6), we defined four groups for a number of selected sequences named *Ornithogalum mosaic virus 1* (OrMV-1) to *Ornithogalum mosaic virus 4* (OrMV4) (Figure 2B). The OrMV groups followed the classification by Gao *et al*. (2018), where OrMV-2 corresponds to Clade A and Clade B is further divided into OrMV groups 1 to 3 (Figure 2B). For both virus species, the subgrouping adhered well to the *Potyviridae* identity demarcation criteria (Supplemental Tables ST5 and ST6).

**Figure 2.**
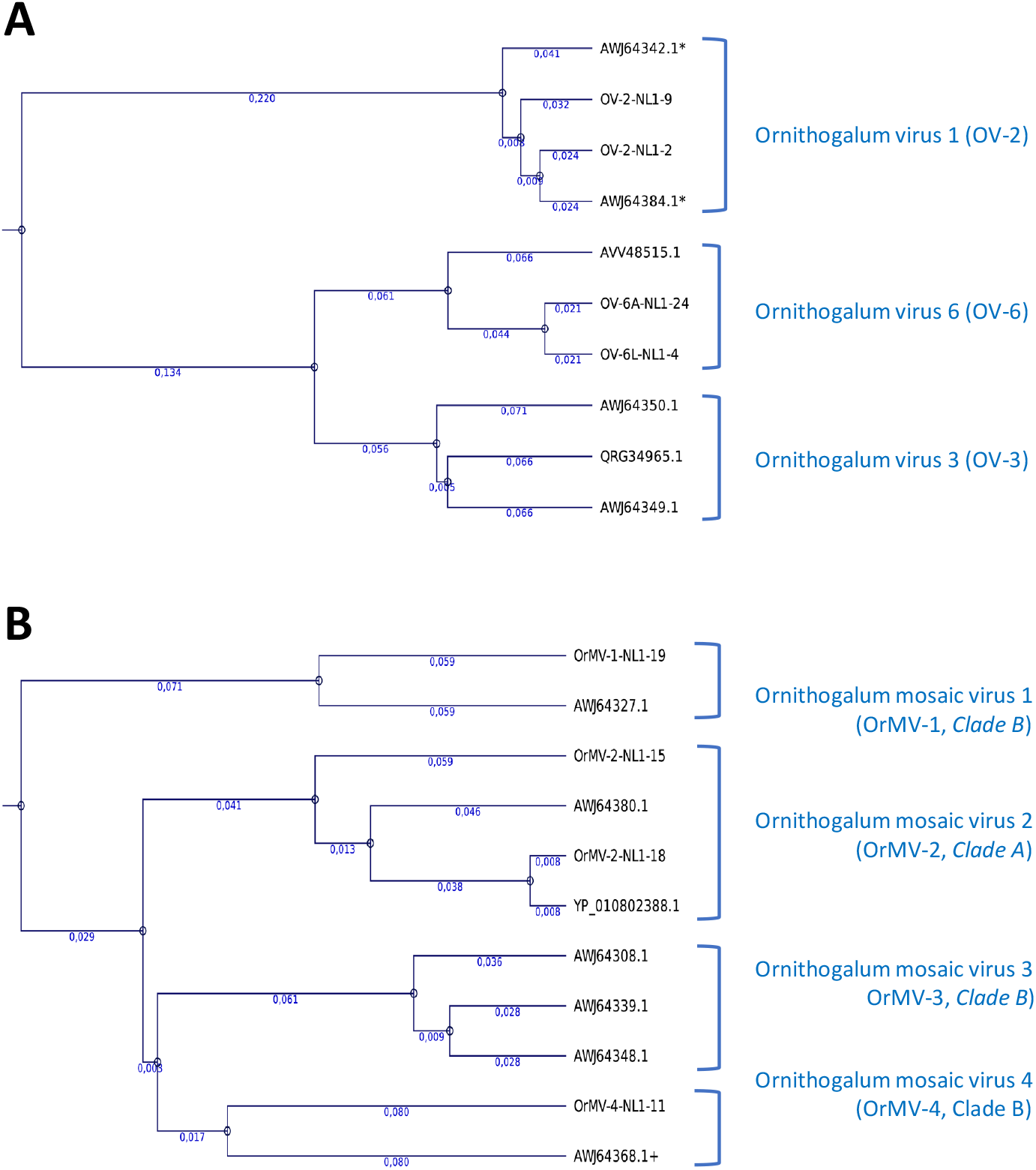
Comparison of similar Ornithogalum virus and Ornithogalum mosaic virus sequences. The Phylogenetic guide trees were generated using the Clustal Omega 1.2.0 algorithm. Branch lengths, indicating the evolutionary distance between sequences are shown in blue below each branch. A) A phylogenetic tree depicting the similarity between selected Ornithogalum virus (OV) polyprotein sequences from GenBank and all new OV variant sequences from this study. The proposed OV classification based on the overall similarity is indicated in blue. B) A phylogenetic tree depicting the similarity between selected Ornithogalum mosaic virus (OrMV) polyprotein sequences from GenBank and all new OrMV variant sequences from this study. The proposed OrMV classification based on the overall similarity is indicated in blue, the clade annotation is from Gao *et al*. (2018).

### Comparing *Ornithogalum* viridae groups

To highlight the differences between these virus groups, we identified and compared the polyprotein cleavage sites for all the selected *Potyviridae* (Table 2). While certain cleavage sites appeared specific to subgroups of the *Ornithogalum* viruses, this specificity was not observed in the *Ornithogalum mosaic* viruses (Table 2). Similarly, the active site of the RNA-dependent-RNA-polymerase (RdRp, NIb) consisting of a 120 bp catalytic domain did not exhibit convincing positions that could serve as markers for virus group classification (Supplemental Figure SF1).

Although comparing polyproteins by identity seemed a solid method for classifying Potyviruses in groups, comparing the mature proteins revealed considerable variation in their identity contrasts (Table 3). Especially, proteins P1 and P3 often differ beyond the demarcation identity values set for the *Potyvirus* polyprotein classification. For example, OrMV-2-NL1-15 shows an identity score of just 60% for P1 and 80% for P3 compared to OrMV-2-NL1-18, despite their polyproteins having an 88% identity (Table 3). A similar pattern is observed for OrMV-NL1-19 and OrMV-1-Awj64327.1 (Table 3).

The conserved interspecies high flexible disorder for the P1, CP, and NIa-VPg proteins as reported by Charon *et al*. (2016) between ten major Potyvirus species, not including the Ornithogalum Potyviruses, only appeared to be reflected in the P1 protein and somewhat in the Cp protein , but not in the NIa-VPg protein of the Ornithogalum Potyviruses. Whereas this appears to be the opposite for the medium disordered P3 protein. Given the various biological functions associated with each mature protein (Charon *et al*., 2016), interpreting these differences remains challenging.

### Overview virus presence

Altogether we encountered viruses from four different families (Table 1 and Supplemental Tables ST3 and ST4). Although the presence of viruses in plants at sampling is best determined by counting virus-related RNAseq reads, the presence of siRNA can reveal current, as well as past virus presence (Malavika *et al*., 2023). Therefore, the (past) presence of viruses in the *Asparagales* plants from the botanic garden was determined by the occurrence of virus-related siRNA. For 13 virus or virus variants, we counted the sRNA-seq reads, presumably representing siRNAs, that completely matched a virus genome sequence (Table 1). The virus-related sRNA-seq read counts ranged from a few hundred to nearly 17 million per sample.

**Table 1.**
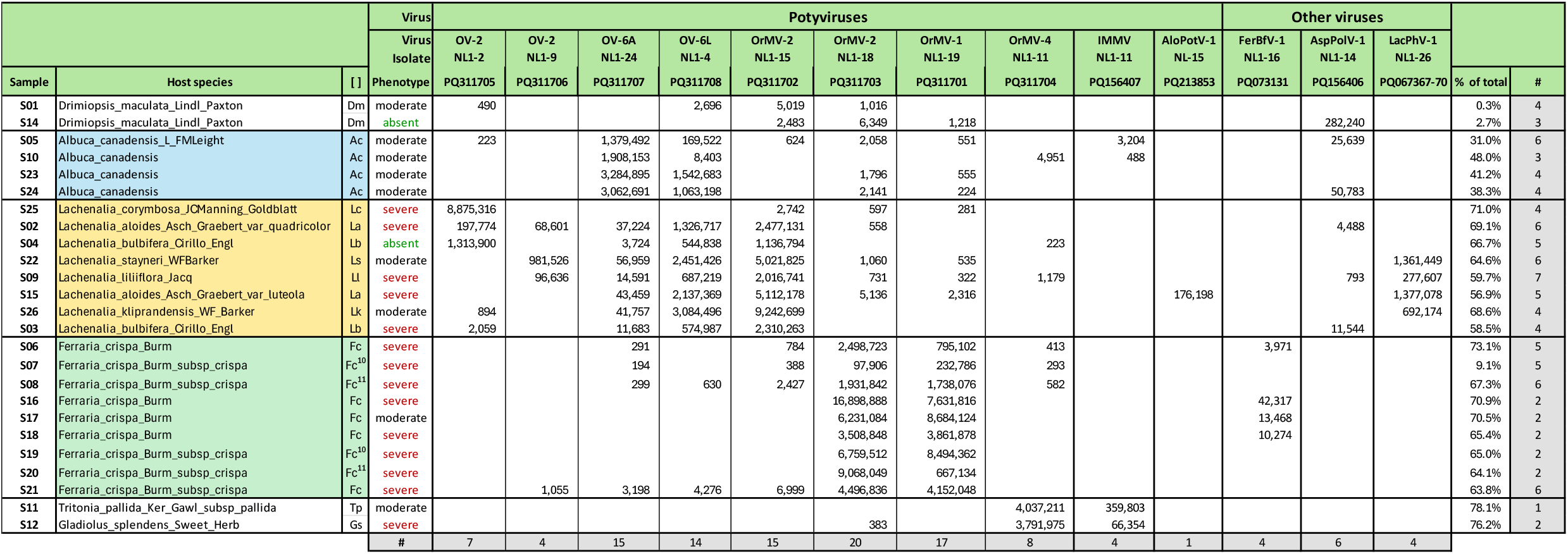
Virus occurrence in Asparagales plants from the Amsterdam botanic garden.

**Table 2.**
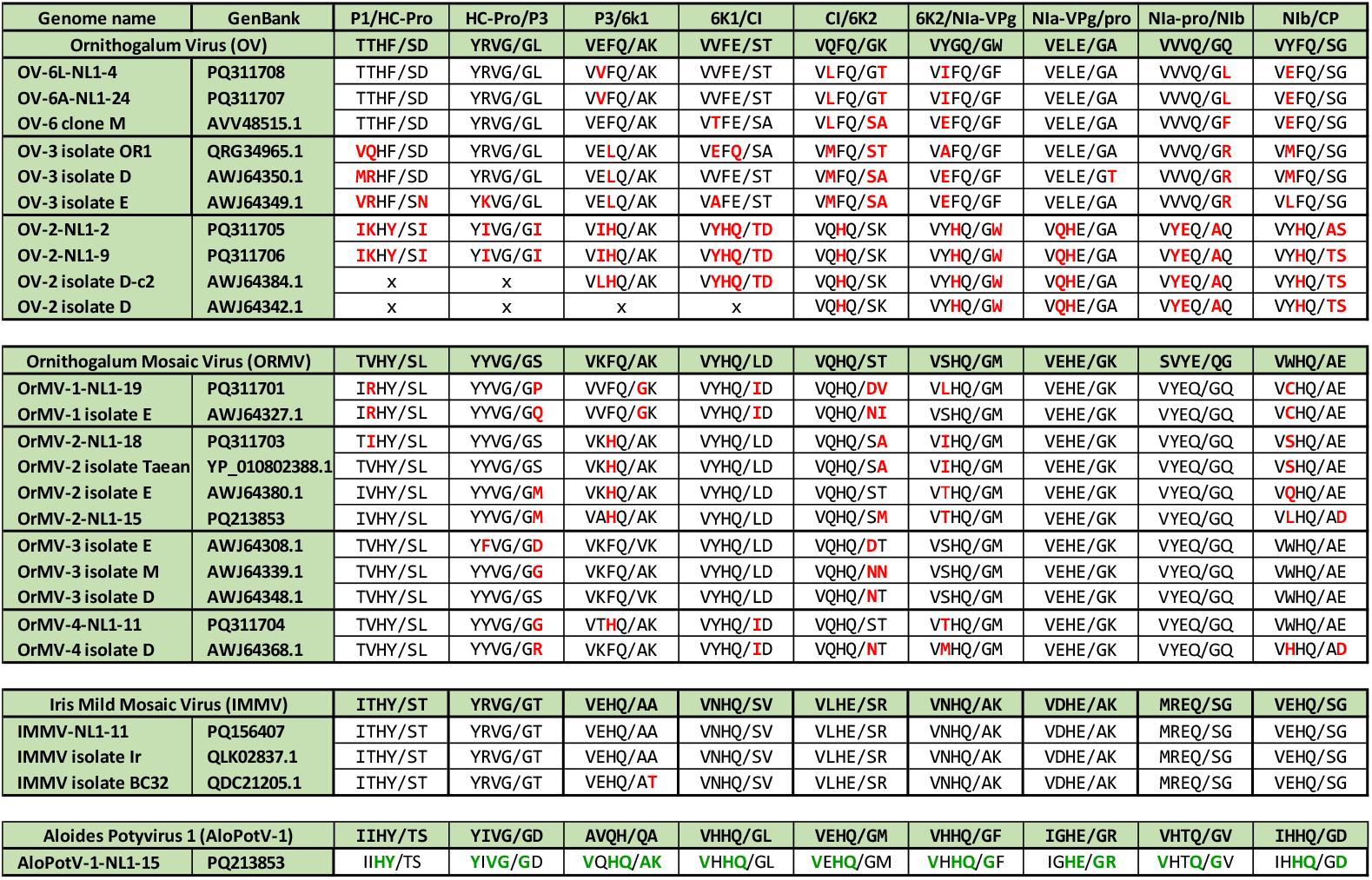
Potyvirus polyprotein cleavage sites.

**Table 3.** Pairwise comparison of the mature protein aa-sequences of one or two selected viruses per Potyvirus subgroup.

| Virus-A | versus Virus B | P1 | HC Pro | P3 | 6K1 | CI | 6K2 | Nla VPg | Nla_pro | N1b | CP | Polyprotein |
| --- | --- | --- | --- | --- | --- | --- | --- | --- | --- | --- | --- | --- |
| OV-2-NL1-2 | OV-2-NL1-9 | 73.4% | 95.0% | 83.1% | 100.0% | 96.2% | 94.3% | 91.8% | 95.1% | 95.4% | 89.1% | 92.8% |
|  | OV-3 AWJ64350.1 | 22.3% | 49.0% | 27.1% | 33.3% | 55.0% | 28.3% | 52.8% | 39.9% | 57.4% | 64.7% | 46.4% |
|  | OV-6 AVV48515.1 | x | 48.4% | 24.9% | x | 53.6% | 30.2% | 49.5% | 40.3% | 55.8% | 58.2% | 47.4% |
| OV-3 AWJ64350.1 | OV-3 QRG34965.1 | 59.7% | 88.9% | 87.0% | 81.1% | 88.2% | 84.9% | 81.9% | 91.8% | 91.3% | 92.9% | 86.2% |
|  | OV-6 AVV48515.1 | 36.0% | 79.5% | 58.0% | 73.6% | 82.3% | 75.5% | 86.2% | 93.4% | 78.7% | 82.1% | 75.9% |
| OV-6 AVV48515.1 | OV-6A-NL1-24 | 58.0% | 91.3% | 80.9% | 83.0% | 93.9% | 86.3% | 86.7% | 95.9% | 90.2% | 90.3% | 86.9% |
|  | OV-6L-NL1-4 | 59.7% | 90.4% | 79.7% | 81.1% | 93.4% | 88.2% | 86.2% | 94.2% | 90.6% | 89.6% | 86.8% |
| OrMV-1-NL1-19 | OrMV-1 AWJ64327.1 | 63.2% | 95.8% | 81.3% | 92.3% | 93.7% | 77.4% | 91.7% | 91.4% | 88.8% | 92.1% | 88.3% |
|  | OrMV-2-NL1-15 | 38.3% | 83.6% | 55.5% | 76.9% | 82.0% | 45.3% | 77.9% | 78.19.0% | 82.1% | 74.0% | 74.0% |
|  | OrMV-2-NL1-18 | 40.6% | 83.8% | 54.3% | 76.9% | 81.5% | 50.9% | 79.7% | 83.8% | 81.1% | 81.9% | 74.4% |
|  | OrMV-3 AWJ64308.1 | 41.1% | 82.9% | 54.6% | 80.4% | 82.7% | 58.5% | 78.6% | 78.6% | 81.7% | 85.8% | 75.0% |
|  | OrMV-4-NL1-11 | 38.6% | 83.1% | 58.9% | 76.9% | 81.9% | 64.7% | 78.1% | 78.6% | 81.7% | 83.4% | 75.2% |
| OrMV-2-NL1-15 | OrMV-2-NL1-18 | 60.4% | 92.8% | 80.2% | 88.5% | 93.9% | 77.4% | 90.6% | 91.3% | 91.3% | 93.7% | 87.9% |
|  | OrMV-3 AWJ64308.1 | 45.5% | 85.3% | 63.4% | 86.5% | 85.2% | 69.4% | 85.9% | 88.4% | 84.4% | 85.3% | 79.3% |
|  | OrMV-4-NL1-11 | 48.5% | 87.1% | 67.5% | 92.3% | 86.7% | 66.0% | 84.9% | 89.7% | 83.4% | 88.5% | 80.6% |
| OrMV-3 AWJ64308.1 | OrMV-3 AWJ64348.1 | 73.3% | 94.5% | 86.2% | 100.0% | 95.1% | 92.5% | 94.3% | 95.0% | 95.8% | 92.9% | 92.0% |
|  | OrMV-4-NL1-11 | 45.9% | 85.3% | 64.7% | 90.4% | 85.0% | 69.8% | 84.4% | 91.3% | 84.4% | 93.7% | 80.2% |
| OrMV-4-NL1-11 | OrMV-4 AWJ64368.1+ | 54.9% | 87.3% | 73.6% | 90.4% | 90.5% | 83.0% | 83.3% | 92.1% | 85.7% | 94.5% | 84.0% |
| IMMV-NL1-11 | IMMV QLK02837.1 | 96.6% | 95.5% | 98.7% | 96.2% | 99.8% | 100.0% | 98.4% | 100.0% | 99.0% | 98.2% | 98.2% |
|  | IMMV_QDC21205.1 | 77.4% | 95.5% | 91.1% | 98.1% | 96.6% | 98.1% | 93.6% | 97.5% | 96.4% | 79.1% | 92.3% |

Identity percentages below the species demarcation threshold of 82% are highlighted in red.

The most obvious observation is that there seems to be quite some (past) virus infections in the analyzed samples, with one sample (S09) containing siRNA related to as much as seven different virus (variants) (Table 1) and none of the samples appeared to be completely virus-free. Conversely, 17 out of 25 samples (68%) contained OV-related siRNA and all samples (100%) contained OrMV-related siRNA (Table 1). From the distribution of observed siRNA sequencing reads, it seems as if the viruses display some preference for certain plant species, OV-2 mostly appeared in *Lachenalia* and *Albuca*, whereas OrMV-1 most highly appeared in *Ferarria crispa* (Table1). Additionally, specific viruses often co-occurred, such as: OrMV-2-NL1-15 and OV6; OrMV-2-NL1-18; and OrMV-1-NL1-19, plus IMMV-NL1-11 and OrMV-4-NL-11. Due to the diverse presence of various virus types and variants it was challenging to identify any correlation between specific viruses and the observed disease phenotypes. It would be interesting to further test the host preference, and the co-occurrence of potyviruses in a larger scale experiment that also includes *Asparagales* plants from other (urban) botanic gardens.

## Concluding remarks

It is widely recognized that only a small portion of the global virome has been discovered. Even though the introduction of high-throughput genome analyses methods (metagenomics) has dramatically accelerated the discovery process, much of the virus research remains concentrated on viruses that impact mammalian health and the health of crops or ornamental plants. There are species for which virome research is just beginning, such as seaweed that promise interesting new virus species (Suttle 2007, Zhao *et al*., 2024). Also, there are many species plant or insect species that inhabit unique environments that harbor specific viruses. One of such unique environments might be urban botanic gardens, which can be likened to islands in their ecological isolation.

Our analysis of 25 *Asparagales* plants from a Dutch urban botanic garden yielded ten new and variant potyviruses, predominantly OV (n = 4) and OrMV (n = 4). The naming of these new and variant potyviruses proved challenging due to the sometimes-confusing nomenclature of viral genome sequences, a complexity introduced by the relatively recent availability of metagenomics. This technology has largely decoupled the genome discovery from the virus phenotypic characteristics and the virus-induced host phenotypes, both of which are crucial for the naming of new viruses. Furthermore, the traditional classification system, which relies on evolutionary relationships and similar phenotypic characteristics might not be suitable for viruses where there may be a genomic continuum.

Aside from the unclear nomenclature, we were not able to link any of the viruses to the observed disease phenotypes of the sampled plants. To establish such links, we need to analyze a larger number of plants and document their phenotypes more accurately. This would be feasible with a variant specific Amplicon-seq approach, based on the sequencing data from this study, allowing us to analyze many relevant plants in this particular hortus botanicus. However, considering the potential high mutation rate of viruses and the fact that Amplicon-seq only will evaluate at best approximately 10% of a potyviral genome, this approach may not be very promising.

In this experiment, we analyzed the plant-derived RNA with both sRNA-seq and RNA-seq. Interestingly we noticed that in some cases siRNA was (abundantly) present while virus RNA was (almost) absent, and vice versa. This suggests past viral infections and recent viral infections, respectively. Regardless, analyzing RNA with both approaches considerably facilitated and improved the discovery of new virus sequences.

Experiments like the one in this study highlight the immense challenge we still face in charting and understanding the global virome and its impact on the health of all living organisms. The discovery of so many new virus variants in a relatively small experiment suggests that (urban) botanic gardens may serve as rich reservoirs of phytoviruses and act as “green islands” that harbor specific phytovirus evolution. More extensive follow-up experiments will provide more insight into these hypotheses.

## Acknowledgments

We would like to express our sincere gratitude to Sarina Veldman, Martin Smit, Iris van Kleinwee, and Reinout Havinga from the Hortus Botanicus in Amsterdam, The Netherlands, for their invaluable support in providing us with plant leaf material for this study. This research was directly and indirectly funded by the Swammerdam Institute for Life Sciences of the University of Amsterdam.

## Data availability

The raw sequence reads have been deposited in the NCBI Sequence Read Archive under BioProject accession number PRJNA1137160. The following virus genome sequences have been deposited in NCBI GenBank: OrMV-1- NL1-19 (PQ311701), OrMV-2-NL1-15 (PQ311702), OrMV-2-NL1-18 (PQ311703), OrMV-4-NL1-11 (PQ311704), OV-2-NL1-2 (PQ311705), OV-2-NL1-9 (PQ311706), OV-6A-NL1-24 (PQ311707), OV-6L-NL1-4 (PQ311708), AloPotV-1 (PQ213853), IMMV-NL1-11(PQ156407).

---

**Supplemental Figure SF1.**
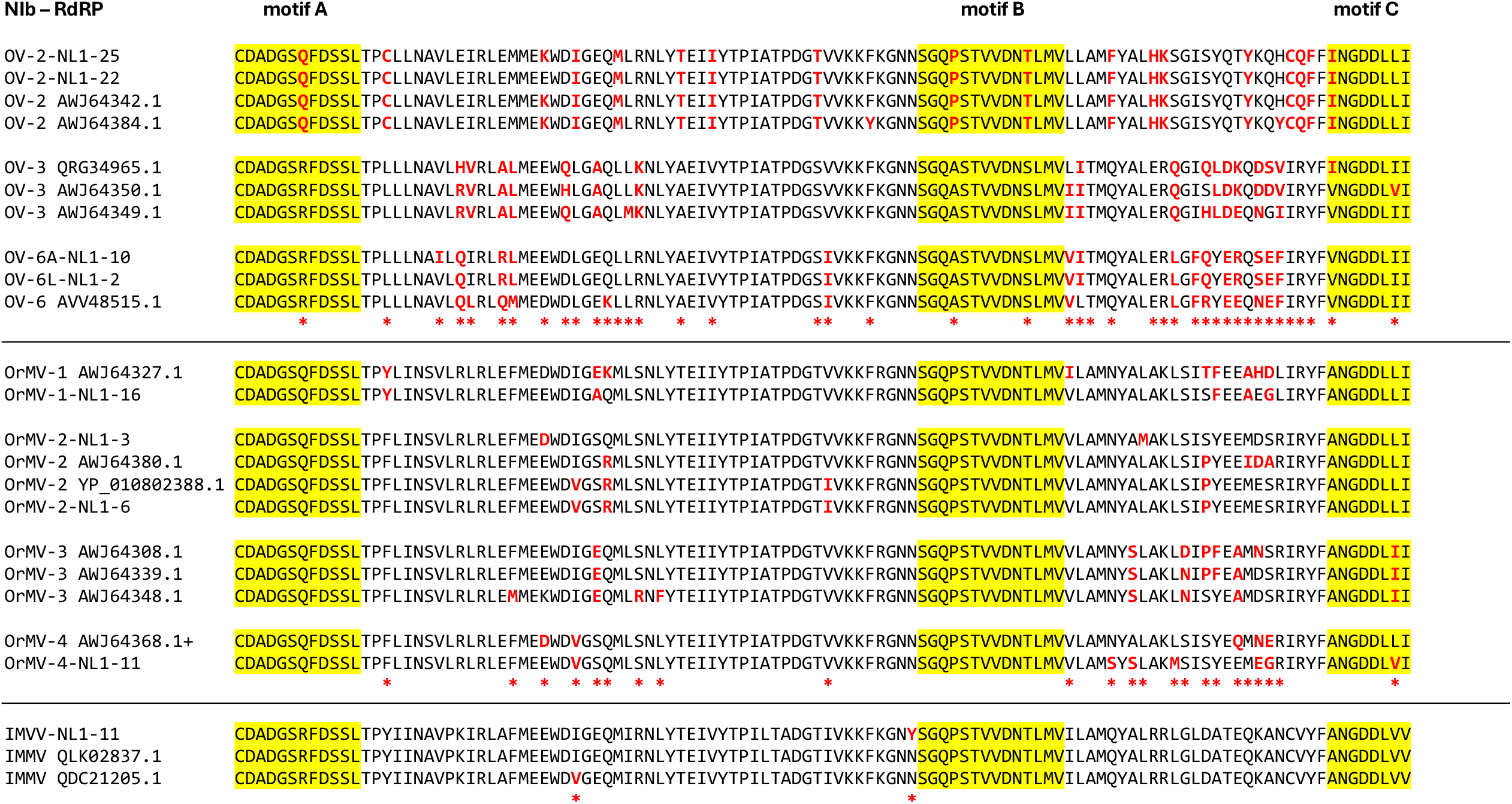
Similarity between RdRp core catalytic motifs A, B, and C (yellow)of the *Ornithogalum* viruses (OV), the *Ornithogalum* mosaic viruses (OrMV), and the Iris mild mosaic viruses (IMMV) as determined by the palmID; viral RdRP (https://serratus.io, Babaian and Edgar, 2022). Red indicates the amino acids that are different from each amino acid prevalent sequence.

**Supplemental Table ST1.** Experiment set-up with sequencing read counts.

| Sample ID | Name | Abbreviation | Pot number | Disease phenotype | Harvest Date | Taxonomy |  |  |  | # of reads/read-pairs |  |
| --- | --- | --- | --- | --- | --- | --- | --- | --- | --- | --- | --- |
|  |  |  |  |  |  | Order | Family | Genus | Species | sRNAseq* | RNAseq |
| S01 | *Drimiopsis maculata* Lindl Paxton | Dm | 2 | moderate | 14_02_2019 | Asparagales | Asparagaceae | *Drimiopsis* | *D. maculata* | 3,570,212 | 34,640,313 |
| S02 | *Lachenalia aloides* Asch Graebert var *quadricolor* | La | 3 | severe | 14_02_2019 | Asparagales | Asparagaceae | *Lachenalia* | *L. aloides* | 5,955,615 | 38,542,291 |
| S03 | *Lachenalia bulbifera* Cirillo Engl | Lb | 4 | severe | 14_02_2019 | Asparagales | Asparagaceae | *Lachenalia* | *L. Bulbifera* | 4,975,216 | 54,405,719 |
| S04 | *Lachenalia bulbifera* Cirillo Engl | Lb | 13 | absent | 14_02_2019 | Asparagales | Asparagaceae | *Lachenalia* | *L. Bulbifera* | 4,498,177 | 48,864,675 |
| S05 | *Albuca canadensis* L FMLeight | Ac | 6 | moderate | 14_02_2019 | Asparagales | Asparagaceae | *Albuca* | *A. canadensis* | 5,094,352 | 74,387,234 |
| S06 | *Ferraria crispa* Burm | FC | 7 | severe | 14_02_2019 | Asparagales | Iridaceae | *Ferraria* | *F. crispa* | 4,511,770 | 39,227,726 |
| S07 | *Ferraria crispa* Burm subsp *crispa* | FC<sup>10</sup> | 10 | severe | 14_02_2019 | Asparagales | Iridaceae | *Ferraria* | *F. crispa* | 3,650,466 | 16,393,962 |
| S08 | *Ferraria crispa* Burm subsp *crispa* | FC<sup>11</sup> | 11 | severe | 14_02_2019 | Asparagales | Iridaceae | *Ferraria* | *F. crispa* | 5,461,449 | 50,399,202 |
| S09 | *Lachenalia liliiflora* Jacq | LI | 14 | severe | 14_02_2019 | Asparagales | Asparagaceae | *Lachenalia* | *L. liliiflora* | 5,193,916 | 41,434,192 |
| S10 | *Albuca canadensis* | Ac | 16 | moderate | 14_02_2019 | Asparagales | Asparagaceae | *Albuca* | *A. canadensis* | 4,000,657 | 31,769,287 |
| S11 | *Tritonia pallida* Ker Gawl subsp *pallida* | Tp | 21 | moderate | 14_02_2019 | Asparagales | Iridaceae | *Tritonia* | *T. pallida* | 5,631,974 | 20,052,549 |
| S12 | *Gladiolus splendens* Sweet Herb | Gs | 22 | severe | 14_02_2019 | Asparagales | Iridaceae | *Gladiolus* | *G. splendens* | 5,061,686 | 36,381,503 |
| S14 | *Drimiopsis maculata* Lindl Paxton | Dm | 1 | absent | 14_02_2019 | Asparagales | Asparagaceae | *Drimiopsis* | *D. maculata* | 10,854,121 | 56,652,319 |
| S15 | *Lachenalia aloides* Asch Graebert var *luteola* | La | 5 | severe | 14_02_2019 | Asparagales | Asparagaceae | *Lachenalia* | *L. aloides* | 15,559,011 | 45,746,374 |
| S16 | *Ferraria crispa* Burm | FC | 7 | severe | 14_02_2019 | Asparagales | Iridaceae | *Ferraria* | *F. crispa* | 34,660,163 | 31,620,407 |
| S17 | *Ferraria crispa* Burm | FC | 8 | moderate | 14_02_2019 | Asparagales | Iridaceae | *Ferraria* | *F. crispa* | 21,189,604 | 22,966,473 |
| S18 | *Ferraria crispa* Burm | FC | 9 | severe | 14_02_2019 | Asparagales | Iridaceae | *Ferraria* | *F. crispa* | 11,280,321 | 32,402,472 |
| S19 | *Ferraria crispa* Burm subsp *crispa* | FC<sup>10</sup> | 10 | severe | 14_02_2019 | Asparagales | Iridaceae | *Ferraria* | *F. crispa* | 23,454,445 | 21,800,891 |
| S20 | *Ferraria crispa* Burm subsp *crispa* | FC<sup>11</sup> | 11 | severe | 14_02_2019 | Asparagales | Iridaceae | *Ferraria* | *F. crispa* | 15,185,230 | 69,625,722 |
| S21 | *Ferraria crispa* Burm subsp *crispa* | FC | 12 | severe | 14_02_2019 | Asparagales | Iridaceae | *Ferraria* | *F. crispa* | 13,570,849 | 21,285,878 |
| S22 | *Lachenalia stayneri* WFBarker | Ls | 15 | moderate | 14_02_2019 | Asparagales | Asparagaceae | *Lachenalia* | *L. stayneri* | 15,290,142 | 37,664,978 |
| S23 | *Albuca canadensis* | Ac | 17 | moderate | 14_02_2019 | Asparagales | Asparagaceae | *Albuca* | *A. canadensis* | 11,732,233 | 24,542,074 |
| S24 | *Albuca canadensis* | Ac | 18 | moderate | 14_02_2019 | Asparagales | Asparagaceae | *Albuca* | *A. canadensis* | 10,911,507 | 49,361,870 |
| S25 | *Lachenalia corymbosa* JCManning Goldblatt | Lc | 19 | severe | 14_02_2019 | Asparagales | Asparagaceae | *Lachenalia* | *L. corymbosa* | 12,504,425 | 35,099,651 |
| S26 | *Lachenalia kliprandensis* WF Barker | Lk | 20 | moderate | 14_02_2019 | Asparagales | Asparagaceae | *Lachenalia* | *L. kliprandensis* | 19,037,058 | 35,044,220 |
|  |  |  |  |  |  |  |  |  |  | 10,913,384 | 38,812,479 |

\* *smallRNA-seq* was done in two separate experiments, which causes the noticeable differences in read counts

**Supplemental Table ST2.** Overview of observed RNA viruses in 25 Asparagales plants from an urban botanic garden.

| Name | [ ] | Capsid | Strand |  | Nucleotide | Genome |  | size | Protein | Taxonomy |  |  |  |
| --- | --- | --- | --- | --- | --- | --- | --- | --- | --- | --- | --- | --- | --- |
|  |  |  |  |  |  | organization |  |  |  | Class | Order | Family | Genus |
| Ornithogalum virus 1-3 | OV1-3 | non-enveloped | positive | ss | RNA | monopartite | linear | ~9.400 | poly | Stelpaviricetes | Patatavirales | Potyviridae | Potyvirus |
| Ornithogalum mosaic virus 1-4 | OrMV1-4 | non-enveloped | positive | ss | RNA | monopartite | linear | ~9.600 | poly | Stelpaviricetes | Patatavirales | Potyviridae | Potyvirus |
| Iris mild mosaic virus | IMMV | non-enveloped | positive | ss | RNA | monopartite | linear | ~9.600 | poly | Stelpaviricetes | Patatavirales | Potyviridae | Potyvirus |
| Aloides Potyvirus 1 | AloPV-1 | non-enveloped | positive | ss | RNA | monopartite | linear | ~9.600 | poly | Stelpaviricetes | Patatavirales | Potyviridae | Potyvirus |
| Ferraria Betaflexivirus 1 * | FerBfV-1 | non-enveloped | positive | ss | RNA | monopartite | linear | ~6,200 | poly | Alsuviricetes | Tymovirales | Betaflexiviridae |  |
| Ornithogalum virus 5 | OV-5 | non-enveloped | positive | ss | RNA | monopartite | linear | 5.885 | 5 | Pisoniviricetes | Sobelivirales | Solemoviridae | Polerovirus |
| Asparagaceae Polerovirus 1 * | AspPolV-1 | non-enveloped | positive | ss | RNA | monopartite | linear | 5.885 | 5 | Pisoniviricetes | Sobelivirales | Solemoviridae | Polerovirus |
| Lachenalia Phenuivirus-1 * | LacPhV-1 | enveloped | negative | ss | RNA | segmented 4 | circular by protein | x | n | Ellioviricetes | Bunyavirales | Phenuiviridae |  |

\* New virus species discovered in this study

**Supplemental Table ST3.** Percentage coverage of virus sequences by siRNA sequencing reads in Asparagales plants from the Amsterdam botanic garden.

|  |  |  | Virus | Potyviruses |  |  |  |  |  |  |  |  |  | Other viruses |  |  |
| --- | --- | --- | --- | --- | --- | --- | --- | --- | --- | --- | --- | --- | --- | --- | --- | --- |
|  |  |  | Virus Isolate | OV-2 NL1-2 | OV-2 NL1-9 | OV-6A NL1-24 | OV-6L NL1-4 | OrMV-2 NL1-15 | OrMV-2 NL1-18 | OrMV-1 NL1-19 | OrMV-4 NL1-11 | IMMV NL1-11 | AloPotV-1 NL-15 | FerBfV-1 NL1-16 | AspPolV-1 NL1-14 | LacPhV-1 NL1-26 |
| Sample | Host species | [ ] | Phenotype | PQ311705 | PQ311706 | PQ311707 | PQ311708 | PQ311702 | PQ311703 | PQ311701 | PQ311704 | PQ156407 | PQ213853 | PQ073131 | PQ156406 | PQ067367-70 |
| S01 | Drimiopsis_maculata_Lindl_Paxton | Dm | moderate | 52.9 | 26.4 | 35.5 | 78.8 | 92.3 | 55.4 | 21 | 14 | 2.5 | 0.9 | 1.8 | 4.0 | 0.0 |
| S14 | Drimiopsis_maculata_Lindl_Paxton | Dm | absent | 3.2 | 0.2 | 5.8 | 59.6 | 89.4 | 89.5 | 78.7 | 0 | 0.7 | 16.6 | 4.0 | 99.7 | 21.5 |
| S05 | Albuca_canadensis_L_FMLeight | Ac | moderate | 32.5 | 5.3 | 99.2 | 94.9 | 59 | 83 | 55.6 | 28.6 | 91.3 | 1.5 | 2.6 | 95.8 | 1.0 |
| S10 | Albuca_canadensis | Ac | moderate | 0.2 | 0 | 99.6 | 38.5 | 3.3 | 24.5 | 15.6 | 94.1 | 52.5 | 0.7 | 1.9 | 0.4 | 0.0 |
| S23 | Albuca_canadensis | Ac | moderate | 2.4 | 0.2 | 99.7 | 97 | 4.5 | 61.1 | 60.5 | 0 | 0.9 | 1.5 | 1.7 | 10.7 | 0.0 |
| S24 | Albuca_canadensis | Ac | moderate | 2 | 0.9 | 99.6 | 97.2 | 12.4 | 32.2 | 36 | 0.7 | 1.9 | 1.1 | 1.7 | 93.4 | 0.0 |
| S25 | Lachenalia_corymbosa_JCManning_Goldblatt | Lc | severe | 99.8 | 17.9 | 2.5 | 20.3 | 45.2 | 43.7 | 42 | 0.7 | 2.7 | 2.5 | 1.4 | 0.0 | 1.4 |
| S02 | Lachenalia_aloides_Asch_Graebert_var_quadricolor | La | severe | 99.1 | 98.7 | 39.2 | 96.2 | 99.3 | 43.7 | 19.8 | 21.2 | 4.9 | 2.4 | 1.4 | 97.1 | 0.0 |
| S04 | Lachenalia_bulbifera_Cirillo_Engl | Lb | absent | 99.5 | 8.2 | 73.1 | 97.7 | 99.4 | 25.8 | 8.5 | 33 | 5.6 | 4.2 | 1.3 | 6.2 | 4.0 |
| S22 | Lachenalia_stayneri_WFBarker | Ls | moderate | 9.2 | 99.5 | 90.2 | 98.4 | 99.6 | 60.3 | 56.3 | 4.7 | 3.8 | 3.2 | 1.7 | 13.1 | 99.8 |
| S09 | Lachenalia_liliiflora_Jacq | Ll | severe | 2.3 | 98.8 | 82.5 | 95.1 | 99.1 | 35.8 | 37 | 77.5 | 18.7 | 2.7 | 1.3 | 76.0 | 97.7 |
| S15 | Lachenalia_aloides_Asch_Graebert_var_luteola | La | severe | 4.9 | 0.7 | 43.8 | 98.2 | 99.6 | 94 | 89.1 | 3.9 | 2.7 | 100.0 | 7.3 | 0.7 | 99.9 |
| S26 | Lachenalia_kliprandensis_WF_Barker | Lk | moderate | 74.1 | 1.1 | 49.7 | 97 | 99.6 | 24.8 | 20 | 4.8 | 3.4 | 3.0 | 1.4 | 0.0 | 99.8 |
| S03 | Lachenalia_bulbifera_Cirillo_Engl | Lb | severe | 86.3 | 5.8 | 37.3 | 96.1 | 99.5 | 14.8 | 6.6 | 27.7 | 4.6 | 2.1 | 1.6 | 99.1 | 0.0 |
| S06 | Ferraria_crispa_Burm | Fc | severe | 4.1 | 1.5 | 38.3 | 16.4 | 35.9 | 99.3 | 99.9 | 42.5 | 7.9 | 2.6 | 95.9 | 1.4 | 0.0 |
| S07 | Ferraria_crispa_Burm_subsp_crispa | Fc<sup>10</sup> | severe | 6.2 | 1.6 | 30.4 | 20 | 47.4 | 97.5 | 99.8 | 40.8 | 5.9 | 1.3 | 1.4 | 0.0 | 0.3 |
| S08 | Ferraria_crispa_Burm_subsp_crispa | Fc<sup>11</sup> | severe | 3.3 | 14 | 40 | 52 | 83.5 | 98.4 | 99.9 | 56.8 | 10.2 | 2.9 | 1.4 | 0.3 | 5.9 |
| S16 | Ferraria_crispa_Burm | Fc | severe | 15.6 | 1.8 | 1.3 | 6.5 | 20.3 | 99.9 | 99.9 | 7.5 | 3.1 | 3.5 | 99.9 | 1.1 | 1.3 |
| S17 | Ferraria_crispa_Burm | Fc | moderate | 5.6 | 1.6 | 0.3 | 2.3 | 13.6 | 99.8 | 99.9 | 6.3 | 3.0 | 3.3 | 99.2 | 0.0 | 0.0 |
| S18 | Ferraria_crispa_Burm | Fc | severe | 3.5 | 0.7 | 2.9 | 1.6 | 11.9 | 99.7 | 99.9 | 4.8 | 2.6 | 3.0 | 98.8 | 0.0 | 0.0 |
| S19 | Ferraria_crispa_Burm_subsp_crispa | Fc<sup>10</sup> | severe | 4.3 | 1.5 | 0.7 | 4.2 | 15.8 | 99 | 99.9 | 4.9 | 3.2 | 3.2 | 1.7 | 0.4 | 0.0 |
| S20 | Ferraria_crispa_Burm_subsp_crispa | Fc<sup>11</sup> | severe | 4.1 | 0.2 | 1.3 | 4.4 | 12.8 | 99.2 | 99.9 | 4.6 | 3.0 | 3.2 | 1.4 | 0.0 | 0.0 |
| S21 | Ferraria_crispa_Burm_subsp_crispa | Fc | severe | 3.6 | 75.4 | 83.8 | 79.9 | 94 | 98.9 | 99.9 | 4.6 | 3.6 | 3.4 | 1.4 | 0.7 | 38.7 |
| S11 | Tritonia_pallida_Ker_Gawl_subsp_pallida | Tp | moderate | 24.5 | 0.4 | 1.4 | 13.3 | 24.8 | 4.6 | 2.9 | 99.7 | 99.9 | 1.8 | 1.2 | 0.0 | 0.0 |
| S12 | Gladiolus_splendens_Sweet_Herb | Gs | severe | 1.2 | 0.4 | 1.4 | 1.5 | 3.5 | 42.4 | 18.5 | 99.6 | 99.8 | 1.6 | 1.6 | 0.0 | 0.0 |

NB: Percentages > 30% are highlighted in yellow

**Supplemental Table ST4.**
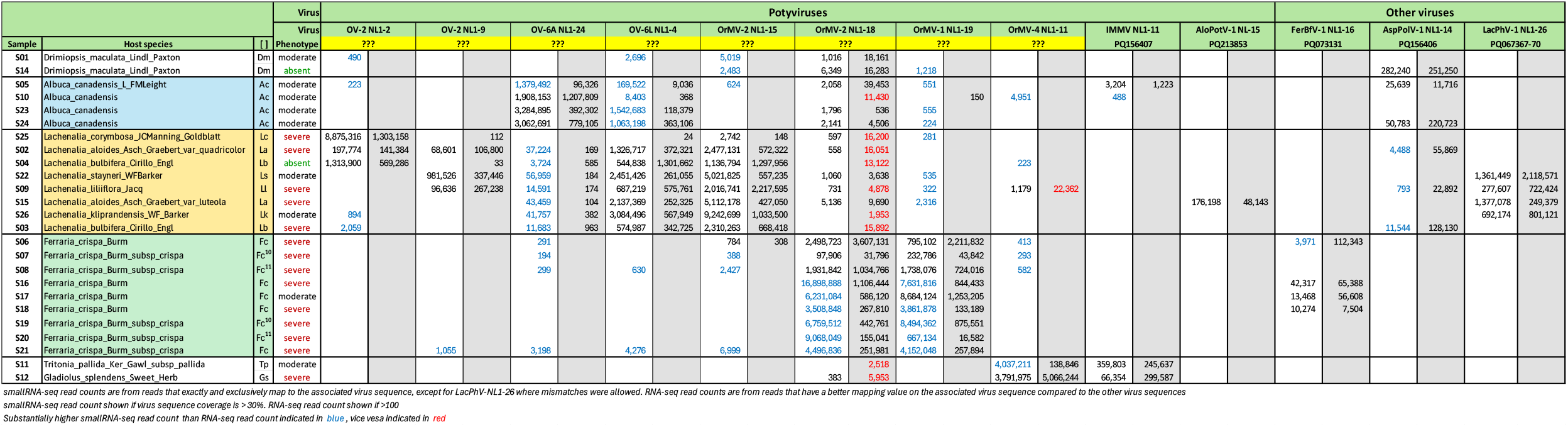
Comparison of virus specific sRNA-seq read count and RNA-seq read counts Asparagales plants from the Amsterdam botanic garden.

**Supplemental Table ST5.** Comparison of Ornithogalum virus polyprotein sequences from representative viruses from GenBank and from variant viruses from this study.

| Type<br>Virus | OV-2 |  |  |  | OV-3 |  |  | OV-6 |  |  |
| --- | --- | --- | --- | --- | --- | --- | --- | --- | --- | --- |
|  | OV-2-NL1-9 | AWJ64342.1* | OV-2-NL1-2 | AWJ64384.1* | AWJ64350.1 | QRG34965.1 | AWJ64349.1 | OV-6A-NL1-24 | OV-6L-NL1-4 | AVV48515.1 |
| OV-2-NL1-9 |  |  |  |  |  |  |  |  |  |  |
| OV-2_AWJ64342.1* | 91.3% |  |  |  |  |  |  |  |  |  |
| OV-2-NL1-2 | 92.8% | 93.1% |  |  |  |  |  |  |  |  |
| OV-2_AWJ64384.1* | 94.2% | 91.7% | 95.3% |  |  |  |  |  |  |  |
| OV-3_AWJ64350.1 | 46.0% | 53.6% | 45.7% | 51.1% |  |  |  |  |  |  |
| OV-3_QRG34965.1 | 4.6% | 52.6% | 45.7% | 51.1% | 86.2% |  |  |  |  |  |
| OV-3_AWJ64349.1 | 45.6% | 52.1% | 45.3% | 50.5% | 85.5% | 86.9% |  |  |  |  |
| OV-6A-NL1-24 | 45.6% | 52.7% | 45.5% | 51.0% | 76.2% | 74.2% | 74.2% |  |  |  |
| OV-6L-NL1-4 | 45.4% | 52.7% | 45.4% | 50.7% | 76.3% | 74.4% | 74.3% | 95.7% |  |  |
| OV-6_AVV48515.1 | 45.6% | 52.8% | 45.6% | 50.9% | 74.8% | 73.8% | 74.4% | 86.9% | 86.8% |  |

Identity percentages below the species demarcation threshold of 82% are highlighted in red. \* = Partial sequence.

**Supplemental Table ST6.** Comparison of Ornithogalum mosaic virus polyprotein sequences from representative viruses from GenBank and from variant viruses from this study.

| Type<br>Virus | OrMV-1 |  | OrMV-2 |  |  |  | OrMV-3 |  |  | OrMV-4 |  |
| --- | --- | --- | --- | --- | --- | --- | --- | --- | --- | --- | --- |
|  | OrMV-1-NL1-19 | AWJ64327.1 | OrMV-2-NL1-15 | AWJ64380.1 | OrMV-2-NL1-18 | YP_010802388.1 | AWJ64308.1 | AWJ64339.1 | AWJ64348.1 | OrMV-4-NL1-11 | AWJ64368.1+ |
| OrMV-1-NL1-19 |  |  |  |  |  |  |  |  |  |  |  |
| OrMV-1_AWJ64327.1 | 88.3% |  |  |  |  |  |  |  |  |  |  |
| OrMV-2-NL1-15 | 74.0% | 75.0% |  |  |  |  |  |  |  |  |  |
| OrMV-2_AWJ64380.1 | 73.6% | 73.6% | 88.5% |  |  |  |  |  |  |  |  |
| OrMV-2-NL1-18 | 73.8% | 74.3% | 87.9% | 90.8% |  |  |  |  |  |  |  |
| OrMV-2_YP_010802388.1 | 74.0% | 74.5% | 88.2% | 90.9% | 98.3% |  |  |  |  |  |  |
| OrMV-3_AWJ64308.1 | 74.3% | 74.2% | 79.4% | 78.7% | 79.4% | 79.6% |  |  |  |  |  |
| OrMV-3_AWJ64339.1 | 74.6% | 74.6% | 79.5% | 78.6% | 79.9% | 79.9% | 93.6% |  |  |  |  |
| OrMV-3_AWJ64348.1 | 74.1% | 74.1% | 79.3% | 78.1% | 79.4% | 79.5% | 92.0% | 94.5% |  |  |  |
| OrMV-4-NL1-11 | 74.6% | 74.7% | 80.6% | 80.4% | 80.7% | 81.1% | 80.2% | 80.3% | 79.8% |  |  |
| OrMV-4_AWJ64368.1+ | 74.8% | 75.2% | 81.2% | 81.0% | 81.5% | 81.6% | 81.0% | 81.4% | 81.3% | 84.0% |  |

Identity percentages below the species demarcation threshold of 82% are highlighted in red.

